# Energy flux controls tetraether lipid cyclization in *Sulfolobus acidocaldarius*

**DOI:** 10.1101/744623

**Authors:** Alice Zhou, Beverly K. Chiu, Yuki Weber, Felix J. Elling, Alec B. Cobban, Ann Pearson, William D. Leavitt

## Abstract

Microorganisms regulate the composition of their membranes in response to environmental cues. Many archaea maintain the fluidity and permeability of their membranes by adjusting the number of cyclic moieties within the cores of their glycerol dibiphytanyl glycerol tetraether (GDGT) lipids. Cyclized GDGTs increase membrane packing and stability, which has been shown to help cells survive shifts in temperature and pH. However, the extent of this cyclization also varies with growth phase and electron acceptor or donor limitation. These observations indicate a relationship between energy metabolism and membrane composition. Here we show that the average degree of GDGT cyclization increases with doubling time in continuous cultures of the thermoacidophile *Sulfolobus acidocaldarius* (DSM 639). This is consistent with the behavior of a mesoneutrophile, *Nitrosopumilus maritimus* SCM1. Together, these results demonstrate that archaeal GDGT distributions can shift in response to electron donor flux and energy availability, independent of pH or temperature. Paleoenvironmental reconstructions based on GDGTs thus capture the energy available to microbes, which encompasses fluctuations in temperature and pH, as well as electron donor and acceptor availability. The ability of Archaea to adjust membrane composition and packing may be an important strategy that enables survival during episodes of energy stress.

**Significance Statement:** Microbial lipid membranes protect and isolate a cell from its environment while regulating the flow of energy and nutrients to metabolic reaction centers within. We demonstrate that membrane lipids change as a function of energy flux using a well-studied archaeon that thrives in acidic hot springs and observe an increase in membrane packing as energy becomes more limited. These observations are consistent with chemostat experiments utilizing a low temperature, neutral pH, marine archaeon. This strategy appears to regulate membrane homeostasis is common across GDGT-producing lineages, demonstrating that diverse taxa adjust membrane composition in response to chronic energy stress.

## INTRODUCTION

Membrane lipids synthesized by microbes from the domain Archaea can be preserved in sediments for millions of years and are used to reconstruct past environmental conditions. Isoprenoid lipids known as GDGTs (glycerol dibiphytanyl glycerol tetraethers) are the main constituents of mono-layer membranes in many archaea (Kates, 1993; Koga and Morii, 2007). These membrane-spanning lipids and can contain up to eight cyclopentane rings in Cren- and Euryarchaeota, or up to four cyclopentane rings with an additional cyclohexane ring in Thaumarchaeota (Sinninghe Damsté *et al.*, 2002) (Figure S1). These ring structures enhance lipid-lipid interactions (Gabriel and Chong, 2000; Nicolas, 2005; Shinoda *et al.*, 2007a; Shinoda *et al.*, 2007b; Pineda De Castro *et al.*, 2016), and ultimately, the packing and stability of the membrane. The relative abundance of these ring structures in GDGTs forms the basis for the widely applied sea- (Schouten *et al.*, 2002; Schouten *et al.*, 2007) and lake- (Powers *et al.*, 2010; Pearson *et al.*, 2011) surface paleotemperature proxy known as TEX_86_.

Membrane-spanning tetraether lipids are the most abundant lipids in many thermophilic archaea (Siliakus *et al.*, 2017) and can make up close to 100% of the membranes of acidophilic archaea (Macalady *et al.*, 2004; Oger and Cario, 2013). GDGTs were first identified in hyperthermophilic archaea isolated from hot springs with average temperatures > 60°C; these lipids were originally interpreted as a primary adaptive feature to high temperatures (De Rosa *et al.*, 1974). Pure culture experiments with thermoacidophilic crenarchaeota show that the number of pentacyclic rings in the biphytanyl chains of GDGTs increases systematically with growth temperature (De Rosa *et al.*, 1980; Uda *et al.*, 2001; Uda *et al.*, 2004). High temperature, however, is not the only challenge to hot spring microbes. Ecosystems that host thermophiles are often characterized by acidity, and archaeal tetraether-based membranes have also been demonstrated to confer tolerance to low pH (Macalady *et al.*, 2004; Boyd *et al.*, 2013). Archaeal membranes composed of tetraethers are relatively impermeable and thus restrict proton influx into the cytoplasm, allowing cells to better maintain homeostasis at low pH and high temperatures (Elferink *et al.*, 1994; Konings *et al.*, 2002). GDGTs may be linked to the survival of Archaea at these environmental extremes, and calditol-linked GDGTs were recently shown to be required for growth of *Sulfolobus acidocaldarius* under highly acidic conditions (pH < 3) (Zeng *et al.*, 2018). Because these lipids are central to the survival of thermoacidophilic archaea, such organisms are ideal targets to study the role of GDGT cyclization.

As a widespread and structurally distinct class of archaeal membrane lipids, GDGTs have been extensively studied with regards to their biophysical properties and role in influencing aggregate membrane behavior. These lipids form stable, highly impermeable monolayers in which individual lipids have low lateral diffusion rates (Jarrel *et al.*, 1998). The incorporation of cyclopentyl rings into these core lipids further increases the thermal stability of membranes. Differential scanning calorimetry experiments on pure lipid films show that thermal transitions are shifted towards higher temperatures as the number of cyclopentane rings increases (Gliozzi *et al.*, 1983). The basis for such trends has been illuminated by molecular dynamics simulations across various timescales both *in vacuo* (Gabriel and Chong, 2000) and in solution (Nicolas, 2005; Shinoda *et al.*, 2007a; Shinoda *et al.*, 2007b; Pineda De Castro *et al.*, 2016). These computational studies demonstrate that membranes comprised of highly cyclized GDGTs are more tightly packed, largely due to the more favorable hydrogen bonding interactions that result from the incorporation of cycloalkyl moieties (Gabriel and Chong, 2000; Shinoda *et al.*, 2007a). Tight packing causes membranes composed of cyclized tetraether lipids to become more rigid than membranes composed entirely of acyclic GDGTs. The extent of membrane packing exerts control on microbial physiology, as membrane fluidity and permeability directly influence how the cell interacts with its environment. The closer packing of GDGTs with more pentacyclic rings can explain why compositional variations in these lipids are directly linked to gradients in temperature (De Rosa *et al.*, 1980; Uda *et al.*, 2001; Uda *et al.*, 2004) and pH (Yamauchi *et al.*, 1993; van de Vossenberg *et al.*, 1998), as well as energy conservation under heat stress (Sollich *et al.*, 2017).

The ability to survive low energy availability may be a defining characteristic of archaea (Valentine, 2007). The relative impermeability of archaeal membranes decreases cellular maintenance energy requirements by reducing inadvertent ion diffusion across the cell membrane (Konings *et al.*, 2002; Hulbert and Else, 2005), implying that the capacity to vary GDGT composition might be an adaptive response to energy limitation. As recently proposed for mesophilic Thaumarchaeota (Hurley *et al.*, 2016), if GDGTs indeed play a role in energy conservation, limiting the energy flux necessary for growth should result in a measurable effect on core GDGT cyclization. We hypothesize this extends to all GDGT-producing archaea, and assess this in a model thermoacidophilic archaeon. In this study we test the effect of limited electron-donor flux on GDGT cyclization in continuous culture (chemostat, Figure S2) experiments with the heterotrophic thermoacidophile *Sulfolobus acidocaldarius* DSM 639. We restrict feed rates of a limiting substrate, sucrose, which acts as both the sole carbon source and electron donor. This strategy allows us to set the specific growth rate of the microbial population while maintaining constant temperature, pH, dissolved oxygen, and chemical composition of the growth medium (Novick and Szilard, 1950; Herbert *et al.*, 1956). This is the first application of continuous culture work to investigate the effects of energy availability on the lipid composition of a thermoacidophilic archaeon. Our chemostat-based approach pares away the confounding variables associated with batch (De Rosa *et al.*, 1980; Uda *et al.*, 2001; Uda *et al.*, 2004, Elling *et al.*, 2014; Elling *et al.*, 2015; Qin *et al.*, 2015; Feyhl-Buska *et al.*, 2016) and mesocosm (Wuchter *et al.*, 2004; Schouten *et al*., 2007) studies, in which metabolic activity and chemical composition change over the course of the experiment and potentially influence lipid distributions.

## RESULTS

### Lipid distributions and ring indices change in response to electron donor supply

Continuous cultures of the crenarchaeon *S. acidocaldarius* produced GDGTs 0-6 at all growth rates, with the average cyclization (expressed as the Ring Index – RI, *see* Methods) increasing at longer doubling times (Figure 1). We observed substantial shifts in the relative abundance of GDGTs with different amounts of cyclopentyl moieties depending on the steady-state growth rates of the cultures (Figure 1). The mean RI changed from 2.04 ± 0.06 at the fastest growth rate to 3.19 ± 0.14 at the slowest growth rate (Figure 1), corresponding to target turnover rates of 9 to 70 h (see Table 1) and inferred doubling times of 7 to 53 h. Changes in RI were caused by shifts in GDGT distributions, as the relative proportion of GDGTs-4, 5, and 6 increased at slower growth rates (Figure 1). The RI at each target turnover rate (Table 1) was significantly different from all other rates at a 90% confidence level (two-sample *t*-test, p < 0.02), except those targets between and 18h and 30h (Figures 2, S2, S4). To test whether collecting biomass over prolonged intervals altered lipid distributions, a direct comparison between cold trap recovery versus instantaneous reactor biomass was carried out in the slowest target turnover rate (70h). Neither GDGT distributions nor RI values differed significantly between biomass collected into cold traps versus biomass sampled directly from bioreactors (Figures 2, S2, S4). Overall, GDGTs were more cyclized at slower reactor turnover times and slower specific growth rates, resulting in higher RI values (Figure 3, S5).

**Figure 1.**
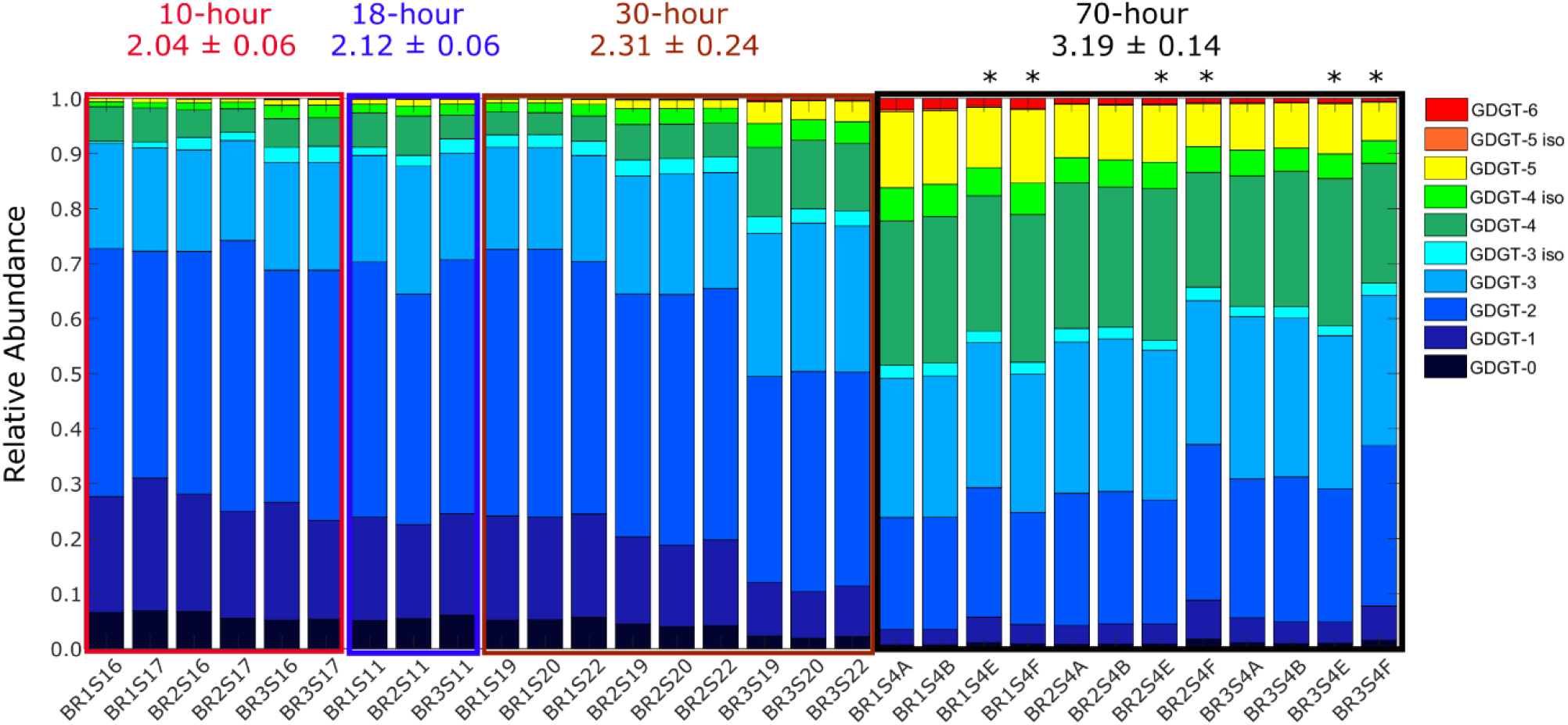
Relative abundances of core GDGTs change solely as a function of doubling time, which we controlled by fixing the provision rate of sucrose to three independent bioreactors (BR1, BR2, BR3). At slower growth rates, lipid distributions are shifted towards more highly cyclized GDGTs-4, 5, and 6. The mean and standard deviations for ring index were averaged across all bioreactors for each target turnover rate, noted above the bar chart. Samples denoted with an asterisk (*) were removed directly from reactors; all other samples were generated from biomass collected into ice-chilled reservoirs.

**Table 1.** Target and mean turnover times for the four dilution rate experiments, as well as the associated inferred population doubling times at steady state. Values account for small volume changes over the course of the experiment. Intervals in which pumping was interrupted are not included.

| Target turnover time (h) | Measured turnover time (h) |  |  | Inferred doubling time (h) |  |  |
| --- | --- | --- | --- | --- | --- | --- |
|  | BR 1 | BR 2 | BR 3 | BR 1 | BR 2 | BR 3 |
| 10 | 9.2 ± 0.7 | 9.9 ± 0.4 | 10.2 ± 0.9 | 6.9 ± 0.7 | 7.1 ± 0.9 | 7.0 ± 0.8 |
| 18 | 17.4 ± 2.3 | 18.5 ± 3.4 | 18.0 ± 4.2 | 12.4 ± 2.1 | 12.2 ± 2.1 | 11.9 ± 3.7 |
| 30 | 29.6 ± 0.4 | 29.9 ± 0.7 | 31.0 ± 1.6 | 21.3 ± 4.8 | 20.4 ± 1.8 | 21.3 ± 1.4 |
| 70 | 70.2 ± 8.7 | 62.2 ± 4.6 | 57.9 ± 4.2 | 51.6 ± 1.9 | 41.3 ± 1.2 | 40.0 ± 1.4 |

**Figure 2.**
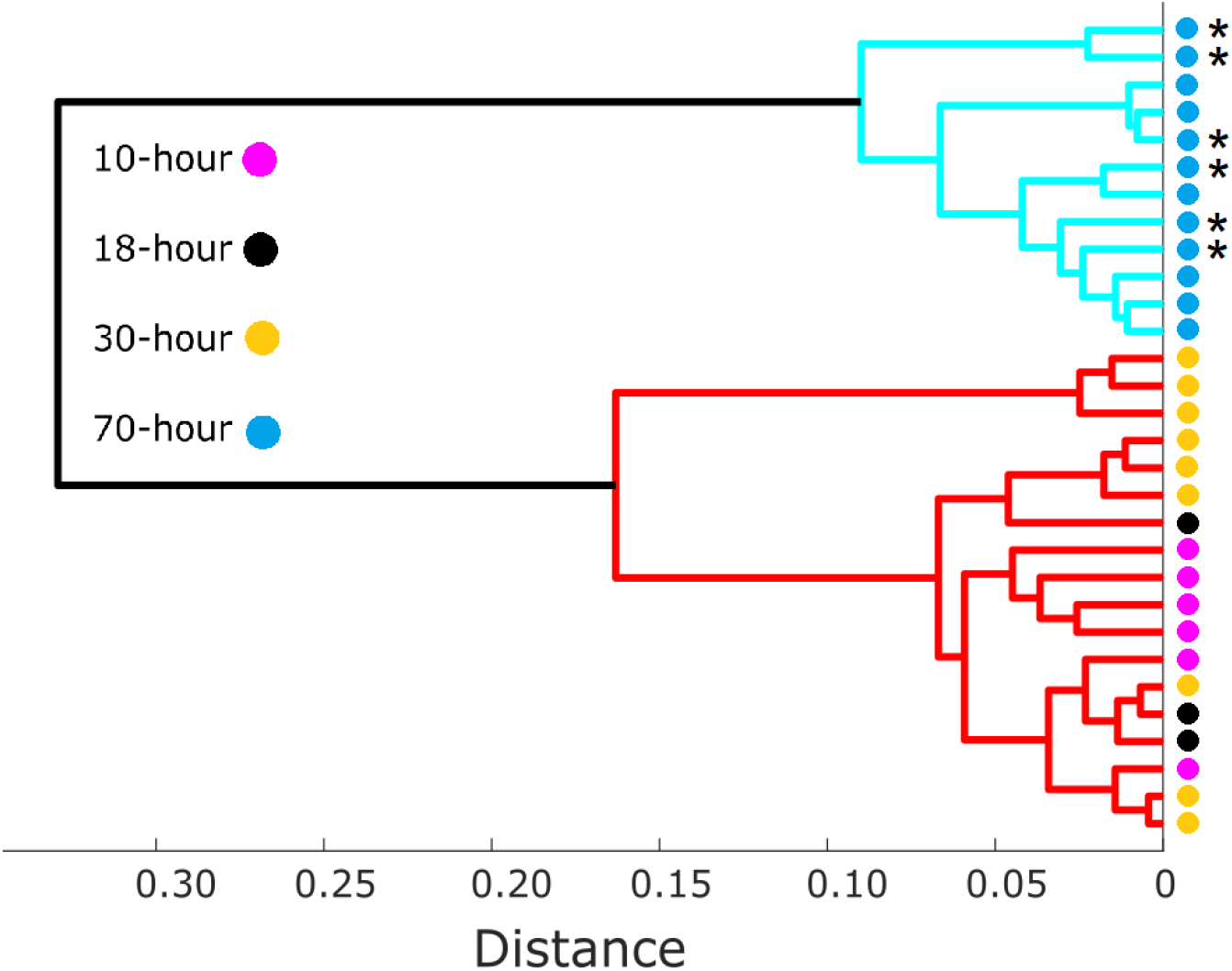
Average linkage dendrogram (cophenetic correlation coefficient = 0.94) showing dissimilarities between normalized GDGT distributions measured in samples collected from cultures spanning 10h to 70h target turnover rate (n = 3). Asterisks (*) are as in Fig. 1.

**Figure 3.**
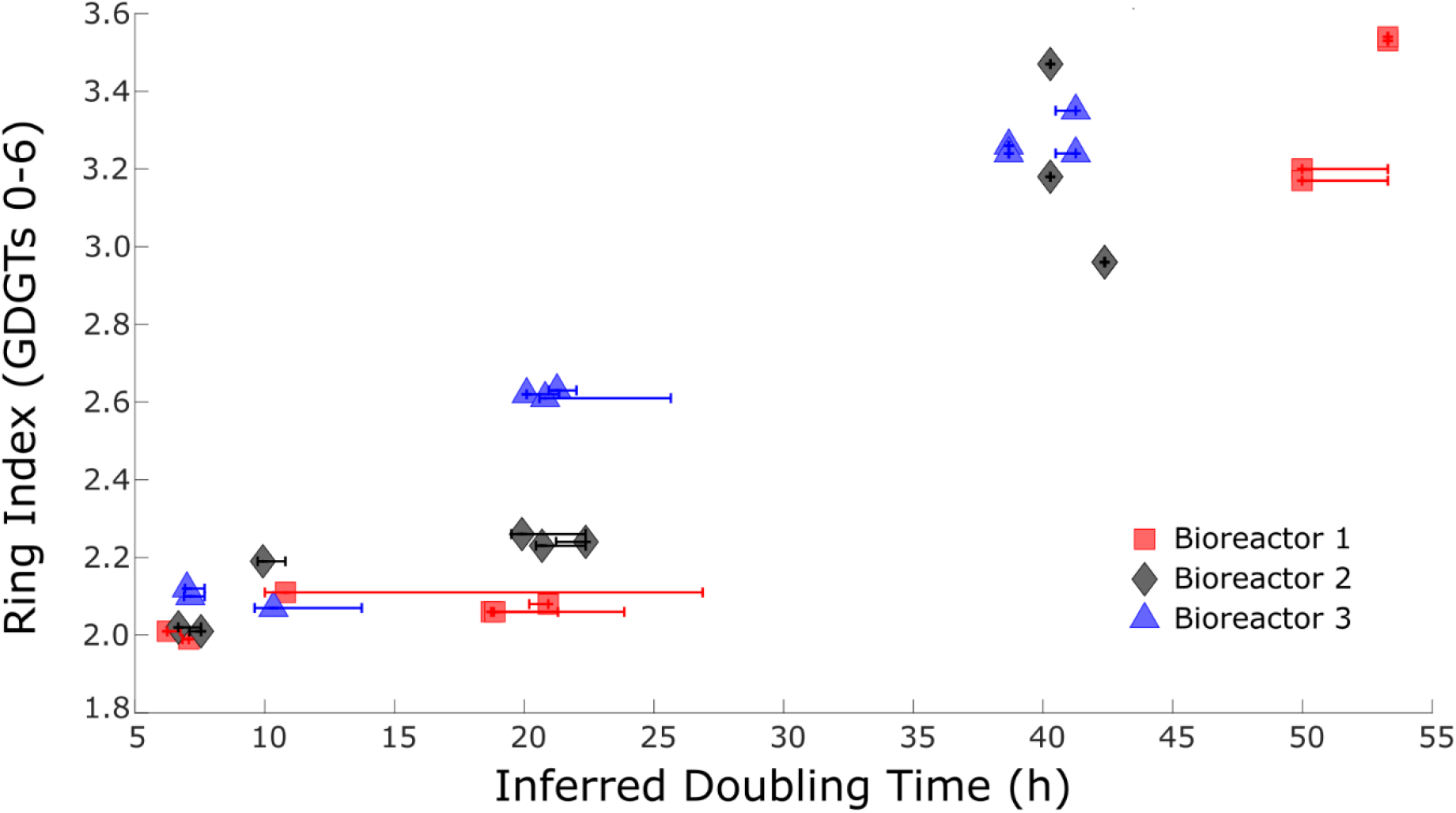
Core GDGT cyclization increased systematically as a function of doubling times in isothermal continuous cultures of *S. acidocaldarius*. Growth rates are controlled by restricting flux of the limiting substrate (sucrose, which serves as both the sole carbon and electron source). X-axis error bars represent the maximum range in inferred doubling times observed during the five reactor turnovers preceding each sampling event.

### Late-eluting GDGT isomers

Small amounts of late-eluting isomers of GDGTs-3, −4, and −5 were observed in all samples (Figures 4 and S6). These molecules are distinct from the early-eluting isomer of GDGT-4 characterized in previous studies (Sinninghe Damsté *et al.*, 2012), but may correspond to minor peaks of putative structural isomers observed in *Sulfolobus solfataricus* GDGTs (Hopmans *et al.*, 2000). The exact structure of these late-eluting isomers remains unknown. Here we identified them based on their retention times and molecular masses in relation to the major GDGTs (Figure S6). We have also observed these late-eluting GDGT isomers in recent batch experiments with *S. acidocaldarius* when the organism was cultivated at conditions associated with the greatest physiological stress, e.g. the highest temperature (80°C) and lowest pH (2.0) tested (data not shown). In this study, the low pH (2.25) may likewise be driving production of late-eluting GDGT isomers. The relative abundances of these isomers were consistent across the entire range of dilution rates tested, which implies that they are not directly involved in a physiological response to energy stress.

**Figure 4.**
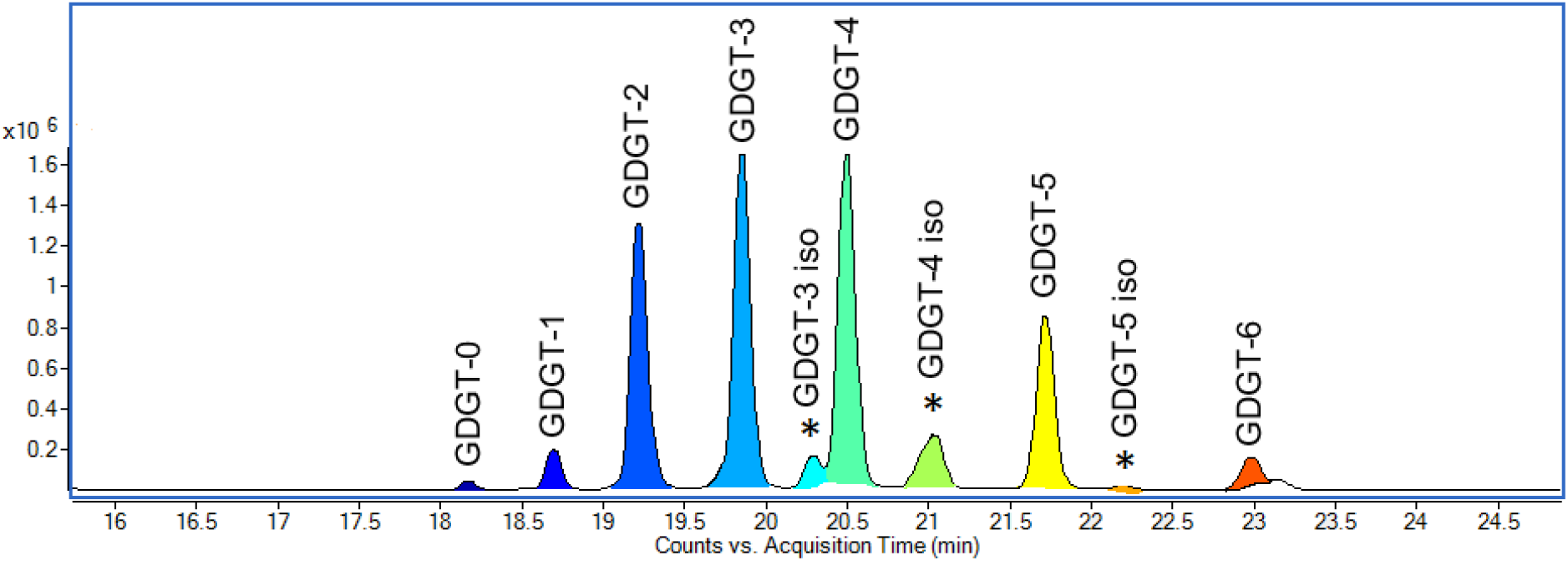
A representative base peak chromatogram of *Sulfolobus acidocaldarius* core GDGTs, obtained through high-performance liquid chromatography/atmospheric pressure chemical ionization mass spectrometry (HPLC/APCI-MS), showing the positions and peak heights of late-eluting isomers (*) relative to major isomers.

## DISCUSSION

Ring index is a measure of the average number of cyclopentane rings within an ensemble of GDGTs. Early laboratory work established an empirical relationship between growth temperature and GDGT cyclization in thermoacidophilic Crenarchaeota (De Rosa *et al.*, 1980; Uda *et al.*, 2001; Uda *et al.*, 2004). Liposome experiments and biophysical models demonstrate that this cyclization is associated with an increase in the degree of membrane packing and a coincident decrease in permeability (Jarrel *et al.*, 1998; Gabriel and Chong, 2000; Nicolas, 2005; Shinoda *et al.*, 2007a; Shinoda *et al.*, 2007b; Pineda De Castro *et al.*, 2016). While RI is a useful summary metric of archaeal GDGT profiles, it provides a non-unique description of a lipid ensemble. For example, a pure GDGT-3 membrane would have the same RI (3.0) as a membrane that is 50% GDGT-1 and 50% GDGT-5. It is unknown whether membranes composed of those example lipid populations would have identical biophysical properties. As such, it may be an oversimplification to draw conclusions based solely on RI values.

The TEX_86_ ratio is a similar metric that applies to GDGTs produced by marine Thaumarchaeota and is also assumed to be driven by changes in growth temperature (Schouten *et al*., 2002). However, recent pure culture experiments with Thaumarchaeota and Crenarchaeota have shown that other factors significantly alter GDGT distributions, TEX_86_, and RI. Both mesophilic Thaumarchaeota and thermo(acido)philic Crenarchaeota appear to use GDGT cyclization to regulate membrane permeability and fluidity in response to a number of environmental stressors (Uda *et al.*, 2001; Macalady *et al.*, 2004; Uda *et al.*, 2004; Elling *et al.*, 2014; Elling *et al.*, 2015; Qin *et al.*, 2015; Siliakus *et al.*, 2017). Specifically, increases in RI during later growth phases in both Crenarchaeota and Thaumarchaeota (Elling *et al.*, 2014; Jensen *et al.*, 2015; Feyhl-Buska *et al.*, 2016), and at reduced O_2_ concentrations in experiments with Thaumarchaeota (Qin *et al.*, 2015), suggest that energy conservation is important in influencing archaeal membrane composition. Since nutrient depletion during later growth phases (in batch cultures) and lower dissolved oxygen results in decreased rates of energy supply, these experiments suggest that there may be a direct feedback between energy availability and cellular membrane composition in GDGT-producing Archaea.

One means to reduce cellular maintenance energy requirements, regardless of surrounding environmental conditions, is to decrease ion permeability across the cytoplasmic membrane (Valentine, 2007). Maintaining a chemiosmotic potential across the membrane is a significant energy expenditure for all living cells, and the spontaneous diffusion of ions across the membrane not through controlled channels (e.g. ATP synthase) is a form of futile ion cycling and imparts a direct energy loss (Hulbert and Else, 2005; Oger and Cario, 2013). More compact membranes decrease the rate at which ion leakage across the membrane can occur, which will lower cellular maintenance energy requirements. Consistent with this idea, RI should increase at lower energy flux, reflecting an increase in membrane packing under energy limitation. Conversely, RI should decrease as energy availability increases, as the cells would tolerate greater ion leakage. The significant increase in the relative abundance of highly cyclized GDGTs at the slowest sucrose supply rate (Figure 3) supports the hypothesis that tighter membrane packing is a response to energy limitation and may indicate a physiological response to low-power environments (c.f. Bradley *et al.*, 2018).

Clear parallels can be drawn between this study and work with another archaeal taxon. Hurley et al. conducted isothermal continuous culture experiments with the mesophilic marine thaumarchaeon *Nitrosopumilus maritimus* SCM1, controlling growth rate by limiting influx of the electron donor, ammonia (Hurley *et al.*, 2016). *N. maritimus* is an ammonia oxidizer of direct relevance to the TEX_86_ paleotemperature proxy, and the chemostat-based approach used by Hurley et al. constitutes the closest experimental analog to this study. Although *S. acidocaldarius* and *N. maritimus* occupy thermally (T > 60 °C vs. T < 30 °C) and chemically (pH < 3 vs. pH > 7) disparate environments and utilize different carbon and energy metabolisms, they both show a positive correlation between specific growth rate and ring index (Figure 5). For both taxa, RI increased with doubling time (Figure 5), although the relative change in average cyclization between end-member growth rates is more pronounced in *S. acidocaldarius* (m = 0.036) than in *N. maritimus* (m = 0.008). This agreement implies that membrane dynamics and cellular bioenergetics are tightly coupled in archaea.

**Figure 5.**
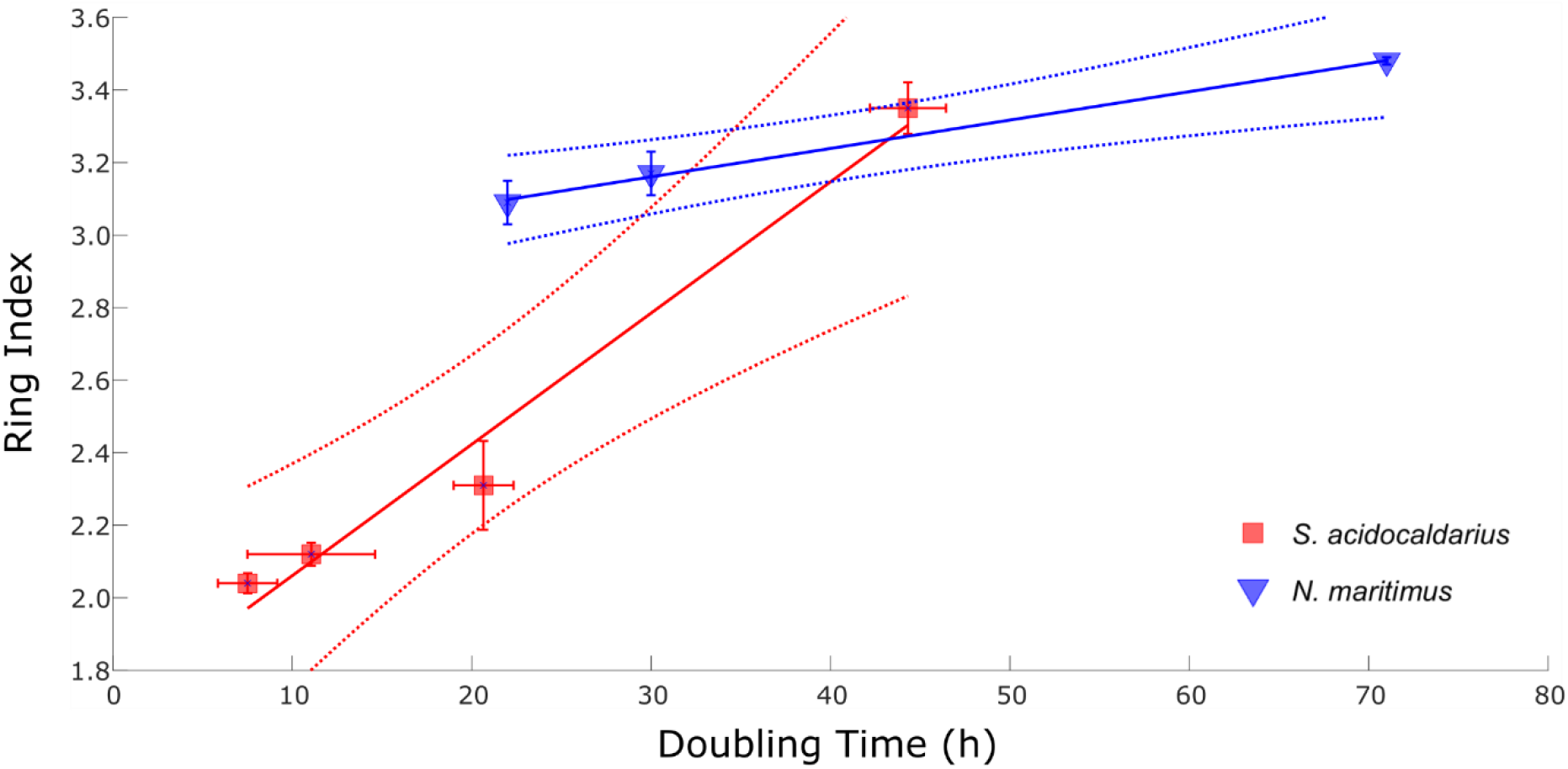
Continuous culture experiments plotting doubling time versus ring index from *S. acidocaldarius* (R^2^ = 0.98; p = 0.01; m = 0.036) relative to *N. maritimus* (R^2^ = 0.99, p = 0.03; m = 0.008) in Hurley et al. (2016). Data from *S. acidocaldarius* are averaged across triplicate bioreactors for each growth rate versus a single reactor for *N. maritimus*. Dashed lines represent 95% confidence intervals.

A mechanistic understanding of how Archaea regulate GDGT distributions is still lacking due to a limited understanding of the GDGT biosynthesis pathway (42). The synthesis of GDGT core lipids involves the highly unsaturated intermediate, di-geranylgeranyl glycerol phosphate (DGGGP). Formation of GDGT-0 from two molecules of DGGGP requires donation of 28 electrons (14 *e*^−^ pairs), perhaps via geranylgeranyl reductase, (GGR) (Nishimura and Eguchi, 2006; Sasaki *et al.*, 2011; Jain *et al.*, 2014). Each ring formed reduces this demand by two electrons, providing a direct physiological link between available energy, the formation of cyclopentyl rings, and thus archaeal membrane lipid composition. The synthesis of ring-containing GDGTs also yields the beneficial outcome of decreased ion permeability. This mechanistic idea explains why cyclization of GDGTs varies with specific growth rate even when temperature and pH are held constant, providing a unifying principle to the environmental factors (temperature, pH, dissolved oxygen) that are observationally correlated with changes in archaeal membrane composition (De Rosa *et al.*, 1980; Gliozzi *et al.*, 1983; van de Vossenberg *et al.*, 1998; Uda *et al.*, 2001; Uda *et al.*, 2004, Boyd *et al.*, 2013; Elling *et al.*, 2015; Siliakus *et al.*, 2017; Sollich *et al.*, 2017).

Changes in GDGT cyclization are more directly quantified by RI than by the TEX_86_ ratio. RI gives the average number of rings of an entire GDGT pool, whereas TEX_86_ expresses the abundance of GDGT-1 relative to the other cyclized GDGTs and was developed specifically to relate lipid distributions to sea surface temperatures (Schouten *et al.*, 2002; Schouten *et al.*, 2007). Ideally, both RI and TEX_86_ would be reported in environmental calibrations or temperature reconstructions, given experimental evidence demonstrating that other factors independently influence GDGT cyclization (Macalady *et al.*, 2004; Elling *et al.*, 2014; Elling *et al.*, 2015; Qin *et al.*, 2015; Feyhl-Buska *et al.*, 2016; Hurley *et al.*, 2016). Preliminary work on the utility of RI as a complementary metric shows that coupling it with TEX_86_ can highlight when GDGT distributions are influenced by non-thermal factors or are behaving differently from average, modern marine communities (Zhang *et al.*, 2016). As expected, RI and TEX_86_ are significantly correlated; deviations of these indices from the modern TEX_86_-RI relationship appear to occur when the GDGT signal is overprinted by the effects of variables that do not predominantly correlate with ocean temperature (Zhang *et al.*, 2016).

Current calibration functions of the TEX_86_ paleotemperature proxy systematically underestimate local SSTs in the tropics and overestimate them at the poles (Tierney, 2014). Given the results of this study and previous work documenting the effect of energy availability on RI and TEX_86_ (Elling *et al.*, 2014; Qin *et al.*, 2015; Hurley *et al.*, 2016), we argue that these biases are likely introduced by temperature-independent environmental and ecological parameters. Marine Thaumarchaeota are ammonia oxidizers whose energy generation is dependent on oxygen as the terminal electron acceptor (Spang *et al.*, 2010 Stahl, D.A. and de la Torre, J.R., 2012). Persistent suboxia and anoxia are common features of modern and ancient restricted basins, such as the proto-Atlantic Ocean during the Cretaceous and Jurassic Oceanic Anoxic Events (Meyer and Kump, 2008). These environmental conditions may directly affect the metabolic activity of marine archaea and therefore impact the TEX_86_ signal in these regions and time intervals. For example, experiments have shown that restricting O_2_ supply in batch cultures can result in as much as a 10°C increase in TEX_86_ temperature estimates in excess of experimental incubation temperature (Qin *et al*., 2015), and a similar shift of 6°C can be induced by altering the electron donor (ammonia) supply in chemostat cultures (Hurley *et al*., 2016). When the results from both distinct experimental approaches are normalized to the same reference frame, biases in TEX86 are explained by changes in growth rate and ammonia oxidation rate (Hurley *et al*., 2016). The impact of energy availability on GDGT distributions is therefore important for interpreting sedimentary records from environments influenced by low O_2_ concentrations or limited electron-donor availability.

Energy limitation may also explain why TEX86-inferred temperatures are anomalously warm (up to +12°C) in modern suboxic settings such as in the permanent oxygen minimum zones (OMZs) of the Eastern Tropical North Pacific Ocean and Arabian Sea, or in seasonally oxygen-deficient regions in coastal upwelling regimes (Basse *et al.*, 2014; Xie *et al.*, 2014; Schouten *et al.*, 2012). In support of this theory, sediments from the Murray Ridge seamount summit, which extends into the Arabian Sea OMZ, yield higher TEX86 temperatures than sediment from adjacent locations that lie below the OMZ (Lengger *et al.*, 2012). Conversely, high energy availabilities are associated with cold biases both in controlled pure culture experiments (Elling *et al.*, 2014; Hurley *et al.*, 2016) and in natural settings such as high-nutrient upwelling systems (Lee *et al.*, 2016). In such instances, high productivity and remineralization rates cause RI- and TEX86-derived temperatures to drop below measured temperatures (Hurley *et al.*, 2018).

Our study establishes that the heterotrophic thermoacidophile *S. acidocaldarius* exhibits a marked membrane-level response to changes in energy availability. These results provide an important experimental counterpart to continuous culture work with the mesophilic and chemoautotrophic archaeon *N. maritimus* (Hurley *et al.*, 2016). In both studies, energy was varied independent of temperature or pH. Together, these experiments suggest that the denser membrane packing associated with core lipid cyclization is an adaptive mechanism that allows diverse GDGT-producing Archaea to cope with energy stress. Quantifying the influence of cellular bioenergetics on archaeal membrane composition may allow for more complete interpretations of GDGT-based records from hot spring, soil, lake, and marine settings. Biomarker-based reconstructions of past environments can be improved by incorporating calibrations that account for local biogeochemical parameters relevant to archaeal energy metabolisms and subsequent membrane reordering.

## METHODS

### Culturing conditions

*Sulfolobus acidocaldarius* DSM 639 was provided by Dr. S-V Albers (University of Freiburg, Germany), and cultured at 70°C in complete Brock medium supplemented with 0.1% NZ-Amine and 0.2% sucrose (see *SI Materials and Methods*) at pH 2.25 (± 0.2). Initial cultures were grown at 70°C and 200 rpm (Innova-42 shaking incubators, Eppendorf). Three 1-L autoclavable glass bioreactors (Applikon, Delft, The Netherlands) were subsequently inoculated with 20 mL of a second-generation culture at mid-exponential phase to an initial optical density of 0.01 at 600 nm (OD_600_).

The three bioreactors were operated in parallel under continuous culture conditions (see *SI Materials and Methods;* Figure S2). Reactors were maintained at 70°C, stirred at 200 rpm (Rushton-type impeller), and aerated with a constant flux of 200 mL/min Zero Air (ultra-high purity, UHP). To reduce evaporative volume loss, excess gas was released through condensers in the reactor headplate. Reactor liquid volume was held constant at 500 mL by equalizing influent and effluent pump rates using a balance control loop (Applikon My-Control software). Temperature, dissolved oxygen, pH, and balance readings were logged continuously using the Lucullus Process Information Management System interface (Applikon).

In four consecutive experiments, the flow rate of the influent and effluent medium was set to target four discrete specific growth rates, μ = 0.069, 0.039, 0.023, 0.010 h^−1^, corresponding to reactor turnover times of T_t_ = 10, 18, 30, and 70 h, respectively. Cell concentrations were monitored by optical density measurements at 600 nm, and coincident fluctuations in dilution rate and optical densities were then used to calculate growth rate and deviation from steady-state.

### Lipid analysis

Biomass collection was initiated after each bioreactor had operated at or within ±10% of steady state for three consecutive turnovers at a given growth rate (Figures S7 and S8). Following collection of effluent into cold traps placed on ice, four 50 mL aliquots from each trap were centrifuged at 3214 x *g* and 4°C for 30 minutes (Eppendorf 5810 R, S-4-104 rotor). The supernatant was decanted after centrifugation and cell pellets were stored at −80°C until ready for freeze-drying and lipid extraction. To determine whether prolonged effluent collection impacted lipid distributions, 10 mL aliquots were also pulled directly from bioreactors during all 70 h experiment sampling events. Aliquots were drawn out through a syringe port and then processed as above.

Core GDGTs were isolated from freeze-dried biomass by acid hydrolysis followed by ultrasonic solvent extraction (e.g. Weber *et al.*, 2017). To this end, freeze-dried cell pellets representing 50 mL aliquots of cell culture were submerged in 3 N methanolic HCl (33% H_2_O) for 3 hours at 70 °C. After cooling, methyl-tert-butyl-ether (MTBE) was added to achieve a MTBE:methanol ratio of 1:1 (vol.) and the samples were agitated using a Qsonica Q500 ultrasonic probe (cup horn, maximum amplitude, 5 minutes total pulse time). Phase separation was induced by changing the solvent composition to MTBE:methanol:hexane (1:1:1, vol.), and the upper organic phase was collected after centrifugation. The total lipid extract (TLE) was subsequently dried under a flow of N_2_ and stored at −20°C in a solution of 1% isopropyl alcohol (IPA) in hexane until analysis.

Core GDGTs were analyzed by ultra-high performance liquid chromatography - atmospheric pressure chemical ionization - mass spectrometry (UHPLC-APCI-MS) using an Agilent 1290 Infinity series UHPLC system coupled to an Agilent 6410 triple-quadrupole mass spectrometer (MS), operated in positive mode (gas temperature: 350 °C; vaporizer temperature: 300°C; gas flow: 6 L min^−1^, nebulizer pressure: 60 psi). Analytical separation of GDGTs was achieved by injecting 2–10 μL of the total lipid extract onto an array of two coupled Acquity BEH HILIC amide columns (2.1 × 150 mm, 1.7 μm particle size, Waters, Eschborn, Germany) maintained at 50°C and fitted with a pre-column of the same material. GDGTs were eluted using a linear gradient from 0.2% to 10% (vol.) IPA in hexane at a flow rate of 0.5 ml/min as previously described (39). At the end of each sample run, the columns were back-flushed with a 70:30 mixture of hexane:IPA (90:10, vol:vol) and IPA:methanol (70:30, vol:vol), and the columns were re-equilibrated to initial condition. The MS was operated in single ion monitoring mode (dwell time 25 ms, fragmentor voltage: 75 V) and GDGTs were quantified by integration of the ion chromatograms of m/z 1302.3 (GDGT-0), m/z 1300.3 (GDGT-1), etc. The ring index (RI) of GDGTs reflects the relative amount of cyclopentyl rings and is defined as:

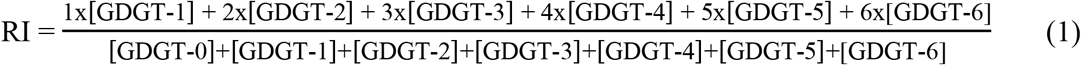

The RI metric compresses GDGT distributions into a single value representing the average number of cyclopentyl rings in GDGTs for each sample.

### Growth rate and doubling time calculations

Cell densities were monitored at regular intervals throughout the experiment by measuring the absorbance at 600 nm (A600) of a 1 mL aliquot pulled directly from each bioreactor. Dilution rates and turnover times were calculated by measuring the volume of effluent pumped out of each reactor per collection interval.

The net rate of change in cell concentration (dx/dt) is a function of the rates of cell division and dilution by sterile medium:

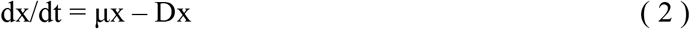

where x = concentration of cells, μ = specific growth rate of the organism, and D = dilution rate (Novick and Szilard, 1950; Herbert *et al.*, 1956). At steady state, the population’s specific growth rate (μ) is equal to the dilution rate (D), such that the concentration of cells does not change with time (dx/dt = 0). Deviation from theoretical steady-state are approximated by calculating the difference between μ and D. Dilution rate is determined by measuring the rate of liquid effluent outflow, and μ is then estimated from measured absorbance data using a rearranged form of Equation 2:

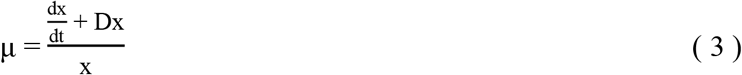

Here dx/dt is the change in A600 between two consecutive time points (i.e. ΔA600/Δt, in h^−1^), D is calculated dilution rate (h^−1^) over the time interval, and x is the measured A600 value at given time point. The value of μ then used to calculate doubling time:

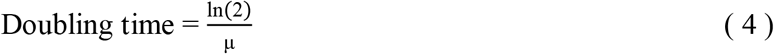

We control and measure both dilution rate and reactor turnover time along with the corresponding biologically relevant metrics, i.e. the calculated specific growth rate and doubling time. At a theoretical steady state in a chemostat, the specific growth rate of a population exactly equals the reactor dilution rate: μ = D. In practice, however, small variances between μ and D can emerge due to the dynamical nature of the reactor system and/or the microbial populations. Deviations from steady state may be driven by transient fluctuations in the internal state of microorganisms, the delivery rate of nutrients, or both. The deviation from theoretical steady-state is expressed in relative terms as:

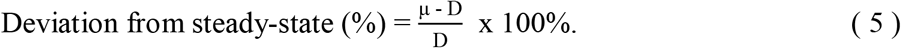

Percent deviation from an idealized steady-state was calculated and recorded each time optical density measurements were taken.

### Statistical analysis

A two-sample *t*-test was used to assess whether the mean RI associated with each growth rate is significantly different from the others (Figure S4). In order to compare the relative abundances of core GDGT variants between samples, we computed a Euclidian distance matrix and visualized dissimilarities between samples by hierarchical clustering using an average linkage technique (Figure S3). All statistical analysis was carried out in MATLAB R2019b; scripts and data files are available online at https://github.com/AliceZhou73/Chemostat-Paper---Code-and-Data-Files.

## Supporting information

SOM Complete

## Acknowledgements

For financial support we thank the ACS-PRF DNI #57209-DNI2 (WDL/YW), the Walter & Constance Burke Fund at Dartmouth College (WDL) and the NASA award NNX15AH79H; Swiss National Science Foundation (YW); and the Gordon and Betty Moore Foundation and NSF-1702262 (AP/FJE). We thank the other members of the Leavitt and Pearson labs for thoughtful discussion and support.

## Supplemental Data

*Code:* https://github.com/AliceZhou73/Chemostat-Paper---Code-and-Data-Files

*Supplementary Information:* Temporary Link: https://figshare.com/s/a8a430482f6e3300c99c

## Notes

https://github.com/AliceZhou73/Chemostat-Paper---Code-and-Data-Files

https://figshare.com/articles/Chemostat_manuscript_dataframe/9640598

