## Supplementary material for "Energy flux controls tetraether lipid cyclization in *Sulfolobus acidocaldarius*": SOM Complete

### SUPPLEMENTAL INFORMATION

#### Materials and Methods

**Medium Preparation.** Concentrated stock solutions of medium components were prepared as described below, with all components dissolved in ultrapure water. To create Complete Brock Medium (CBM), concentrated stocks were combined in the proportions specified and brought up to volume with ultrapure water. The pH of CBM was adjusted to 2.25 using concentrated sulfuric acid ($H_2SO_4$). At the start of this continuous culture experiment, 500 mL CBM was transferred to each bioreactor and autoclaved inside the sealed reactor vessel. Preparation of feed medium used throughout the experiment was conducted in the same manner but scaled up in volume. In most instances, 5 to 6 L of CBM were prepared at a time for each bioreactor and autoclaved directly in 10 L feed bottles. Feed medium was stirred continuously using a magnetic stir bar and fed into bioreactors through a peristaltic pump.

*Complete Brock Medium* was prepared by mixing concentrated stock solutions in the specified volumes. To prepare one liter of CBM: Brock I (1 mL), Brock II/III (10 mL), Fe-solution (1 mL), NZ-Amine solution (5 mL), sucrose solution (10 mL), ultrapure water (973 mL). The pH of this medium was adjusted to 2.25 using concentrated (~9 M) $H_2SO_4$, and medium was autoclaved prior to inoculation.

Brock I (1000x): Concentrated Brock I solution was created by dissolving $CaCl_2$ x $2H_2O$ in ultrapure water to a final concentration of 70 g/L. The solution was autoclave-sterilized at 121°C for 25 minutes.

Brock II/III (100x): Concentrated Brock II/III solution was prepared by dissolving the following components in ultrapure water: $(NH_4)_2SO_4$ (130 g/L), $MgSO_4$ x $7H_2O$ (25 g/L), $KH_2PO_4$ (28 g/L). 50 mL of 2000x trace element solution was added per liter of Brock II/III solution. The mixture was acidified with 1 N $H_2SO_4$ (2 mL/L), and then autoclave-sterilized at 121°C for 25 minutes.

Trace element solution (2000x): The trace element solution was prepared by dissolving the following components in ultrapure water: $MnCl_2$ x $4H_2O$ (3.60 g/L), $Na_2B_4O_7$ x $10H_2O$ (9.0 g/L), $ZnSO_4$ x $7H_2O$ (0.44 g/L), $CuCl_2$ x $2H_2O$ (0.10 g/L), $NaMoO_4$ x $2H_2O$ (0.06 g/L) $VOSO_4$ x $2H_2O$ (0.06 g/L), $CoSO_4$ x $7H_2O$ (0.02 g/L). Note that $Na_2B_4O_7$ x $10H_2O$ is added in excess, which will cause a suspension to form. As such, concentrated trace element solution should be mixed completely before use. This solution was autoclave-sterilized at 121°C for 25 minutes.

Fe-solution (1000x): Fe-solution was prepared by dissolving $FeCl_3$ x $6H_2O$ in ultrapure water to a final concentration of 20 g/L. After complete dissolution, the solution was filter-sterilized through a 0.22 µm pore syringe-driven unit.

NZ-Amine solution (20% w/v): NZ-Amine solution was prepared by dissolving NZ-Amine A in ultrapure water to a final concentration of 20g/100 mL. This solution was autoclave-sterilized at 121°C for 25 minutes.

Sucrose solution (20% w/v): Sucrose solution was prepared by dissolving D-sucrose in ultrapure water to a final concentration of 20g/100 mL. This solution was filter-sterilized using a 0.22 µm pore syringe-driven unit.

**Cultivation.** Three 0.5-L continuous cultures of *Sulfolobus acidocaldarius* DSM 639 were grown aerobically at 70°C in Brock medium (0.1% NZ-Amine, 0.2% sucrose, pH = 2.25). Each culture vessel contained 500 mL of sterile medium inoculated to an initial optical density of 0.01 at 600 nm with 20 mL of a stirred *S. acidocaldarius* batch culture in mid-exponential phase. The chemostat cultures were continuously stirred at 200 RPM via impeller motor and aerated with Zero Air (~20.9% $O_2$) at a constant flux of 200 mL/min. A heating jacket connected to a recirculating cooler system (VWR Heated/Refrigerated Circulator, Model 1196D) was used to regulate temperature while minimizing evaporative loss of reactor contents. Cultures were allowed to reach mid-exponential growth phase (optical density = 0.5) without pumping. After reaching this target optical density, both inflow and outflow pumping were initiated. A peristaltic pump delivered sterile, continuously stirred feed medium into each culture vessel and removed bioreactor contents at the same rate into a sterile collection bottle. Temperature, pH, and dissolved oxygen readings were taken continuously using Applisense BioBundle probes linked to the Lucullus Process Information Management System (PIMS) interface. Cell density was monitored using absorbance spectroscopy at 600 nm using a Genesys 10S UV-Vis spectrophotometer (Thermo Scientific). For these measurements, 1 mL aliquots were sampled every 4-15 h from all bioreactors using a syringe plumbed into the sampling port.

### Supplemental Figures & Tables:

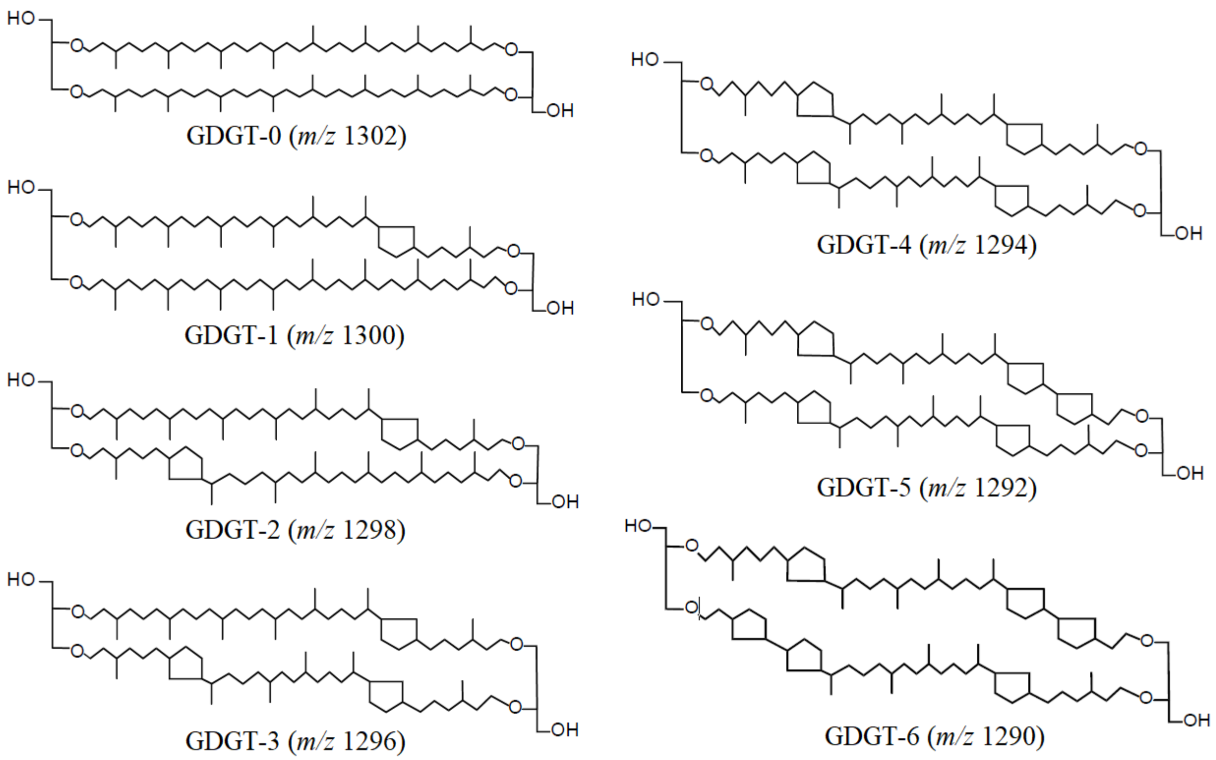


**Figure S1.** Core GDGT lipid structures containing 0 to 6 pentacyclic rings produced by *Sulfolobus acidocaldarius* and observed in this study. Late-eluting isomers were observed for GDGT-3, GDGT-4, and GDGT-5. The stereochemistries of these novel isomers have not yet been resolved.

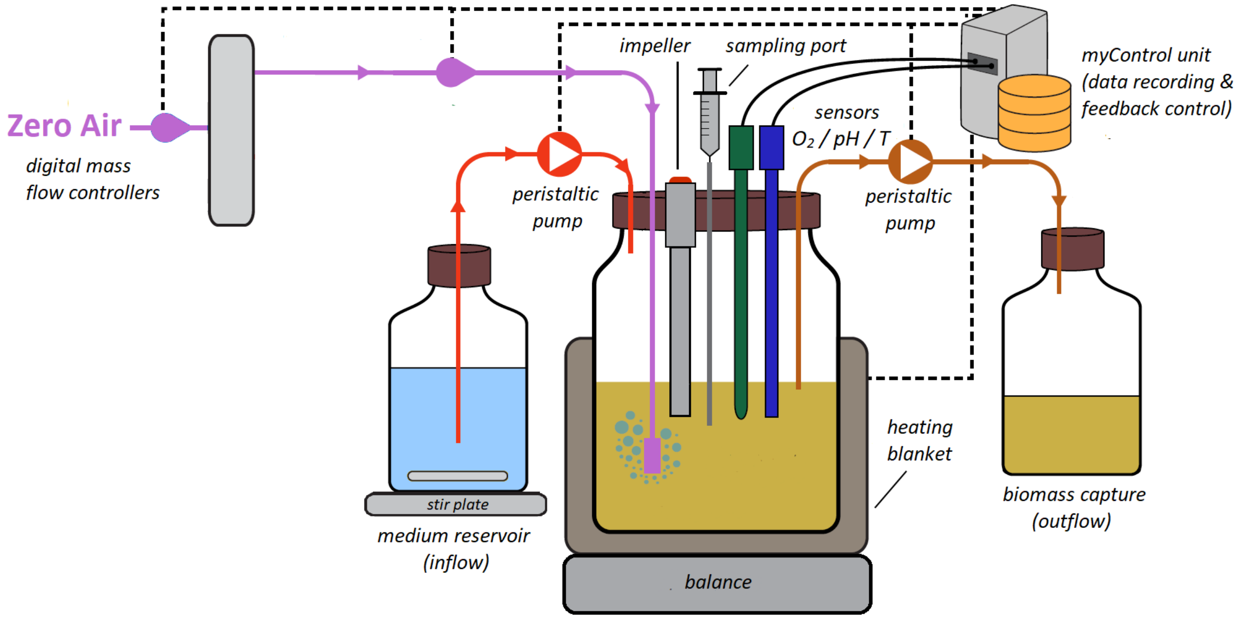


**Figure S2.** Schematic of chemostat setup used in this study (adapted from Sebastian Kopf with permission). The systems were modified from standard Applikon MiniBioBundle 1L glass culture vessels. Each reactor was filled to a total volume of 500 mL and heated to 70°C using a heating jacket and condenser system. The culture was stirred using an impeller motor. Fresh, stirred sterile medium was delivered through a peristaltic pump while reactor contents were pumped out at the same rate into a collection bottle. The reactor volume was sparged with a constant flux of Zero Air (200 mL/min); probes were used to continuously monitor temperature, pH, and dissolved oxygen concentrations over the course of the experiment. Three parallel chemostats were run simultaneously for all experiments, each with its own medium reservoir and control unit.

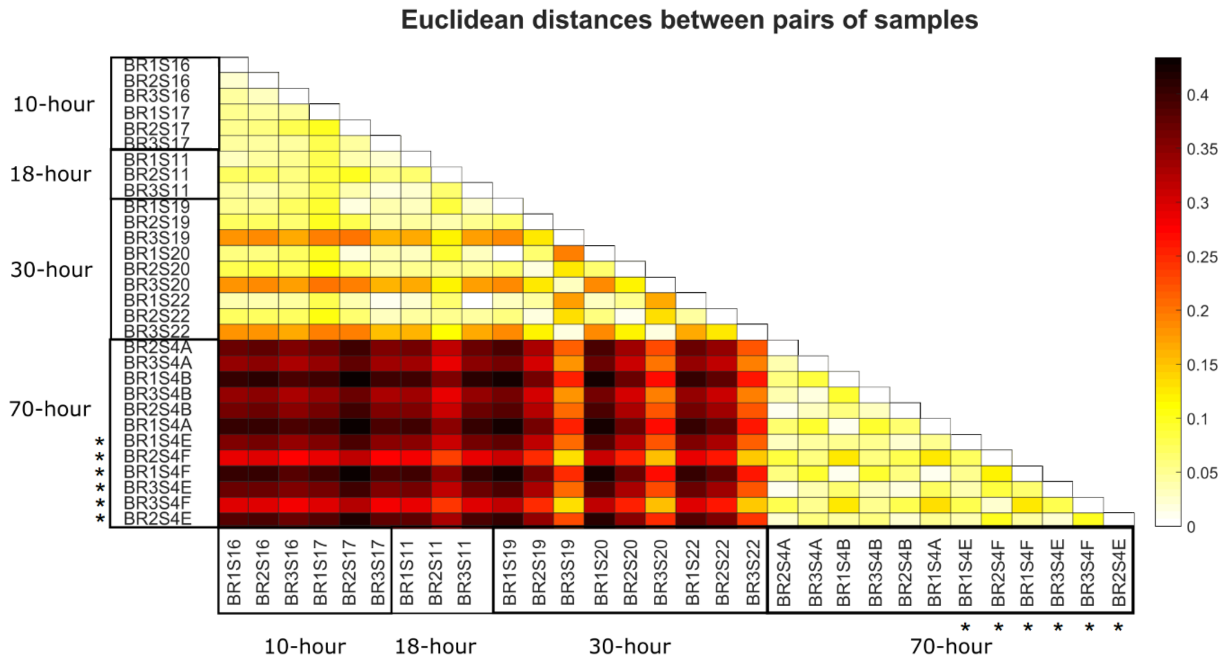


**Figure S3.** A heatmap of Euclidean distances (see Methods for more detailed description of coordinate space) between all pairs of chemostat samples analyzed. Larger distances reflect a greater dissimilarity in the composition of GDGTs between two biomass samples. Samples denoted with an asterisk (*) correspond to biomass pulled directly from reactors; all other data was generated from time-integrated biomass collected in chilled effluent traps.

| Target Turnover Time | 10 h (n = 6) | 18 h (n = 3) | 30 h (n = 9) | 70 h (n = 6) | 70 h, rctr (n = 6) |
|---|---|---|---|---|---|
| **10 h (n = 6)** | | | | | |
| **18 h (n = 3)** | **p = 0.0792**<br>CI=<br>[-0.1757,<br>-0.0124]<br>df = 7 | | | | |
| **30 h (n = 9)** | **p = 0.0216**<br>CI =<br>[-0.4905,<br>-0.0462]<br>df: 13 | **p = 0.2335**<br>CI =<br>[-0.5147,<br>0.1414]<br>df: 10 | | | |
| **70 h (n = 6)** | **p = 1.3191e-09**<br>CI =<br>[-1.3143,<br>-1.0624]<br>df: 10 | **p = 2.3513e-06**<br>CI =<br>[-1.2948,<br>-0.9185]<br>df: 7 | **p = 1.3057e-06**<br>CI =<br>[-1.1566,<br>-0.6834]<br>df: 13 | | |
| **70 h, reactor (n = 6)** | **p = 1.5662e-08**<br>CI =<br>[-1.2560,<br>-0.9540]<br>df: 10 | **p = 1.4596e-05**<br>CI =<br>[-1.2517,<br>-0.7949]<br>df: 7 | **p = 5.4337e-06**<br>CI =<br>[-1.0820,<br>-0.5914]<br>df: 13 | **p = 0.3364**<br>CI =<br>[-0.1006,<br>0.2672]<br>df: 10 | |

**Figure S4.** Pairwise comparisons of mean RI values across turnover time experiments using two-sample *t*-tests. Data is pooled across all reactors for each experimental condition; gold shading represents instances in which the mean RI values associated with turnover times are significantly different ($p < 0.05$). Also reported are the confidence intervals (CI) for the difference in population means between conditions and the degrees of freedom of each *t*-test (df).

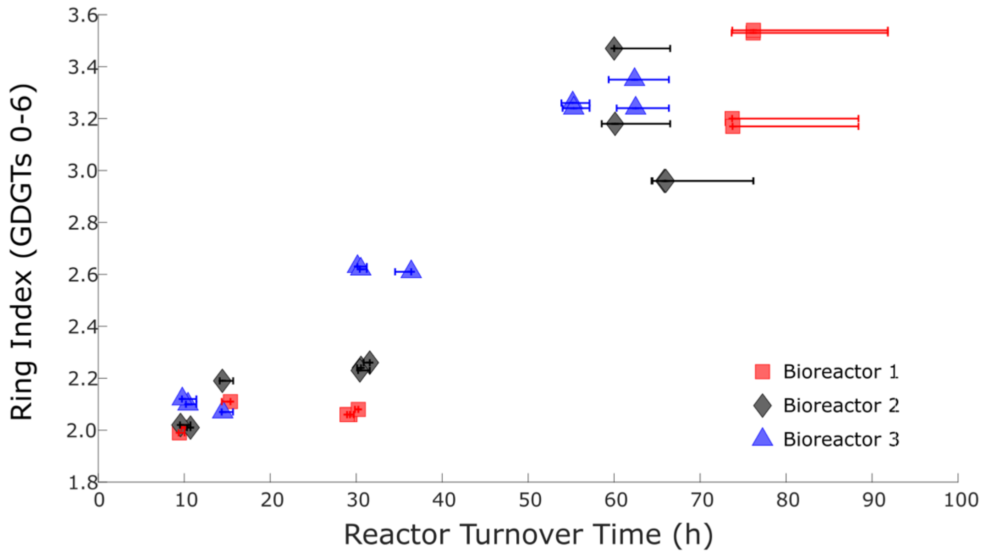


**Figure S5.** Core GDGT cyclization increases systematically as a function of reactor turnover time in isothermal continuous cultures of *S. acidocaldarius*. X-axis error bars represent the range in reactor turnover times observed during the five reactor turnovers preceding each sampling event.

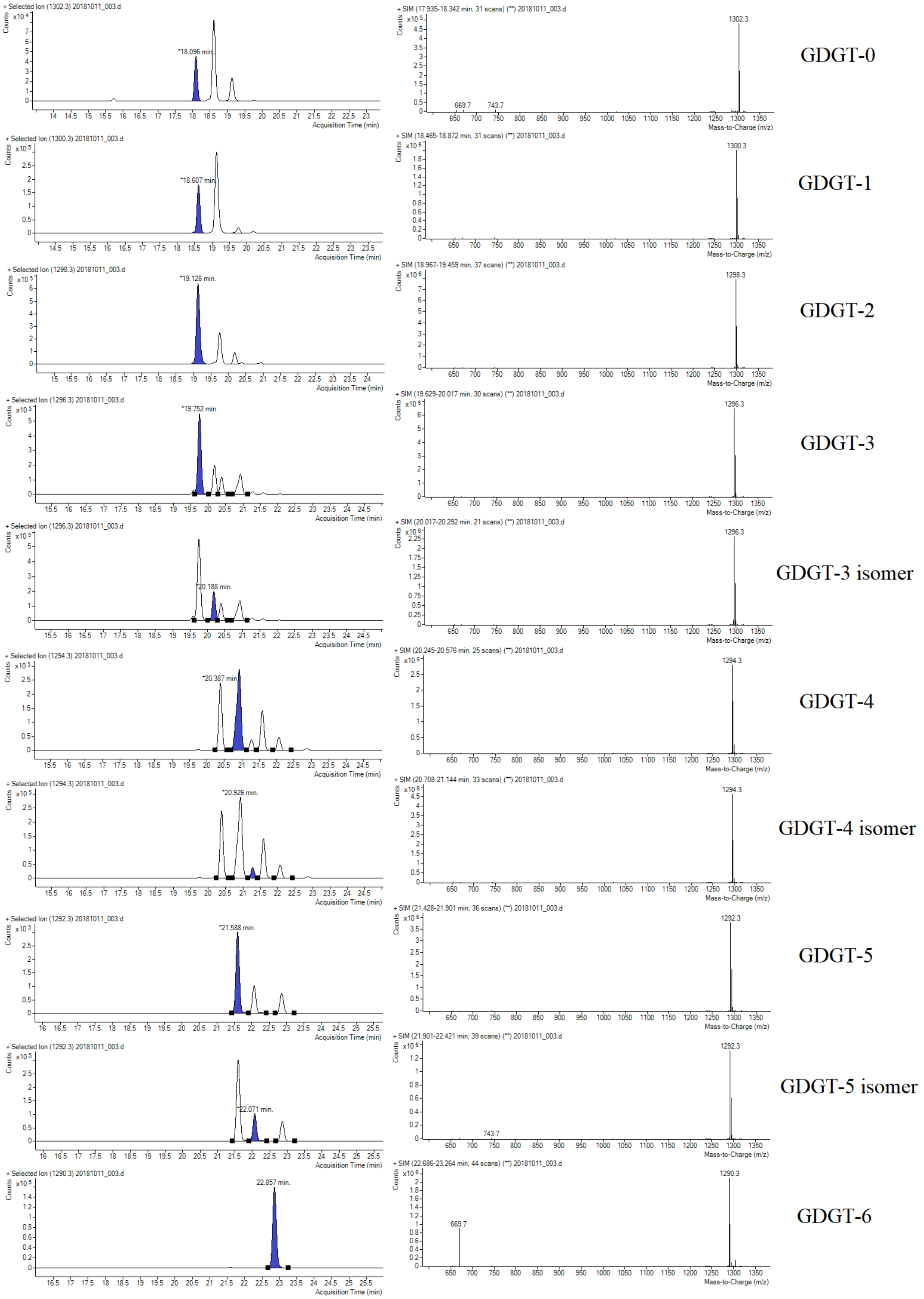


**Figure S6.** LC-MS analyses of *Sulfolobus acidocaldarius* core GDGTs. (A) Total-ion current traces of a representative sample containing GDGTs 0-6. (B) Corresponding atmospheric pressure chemical ionization (APCI) mass spectra of selected GDGT species (shaded peaks).

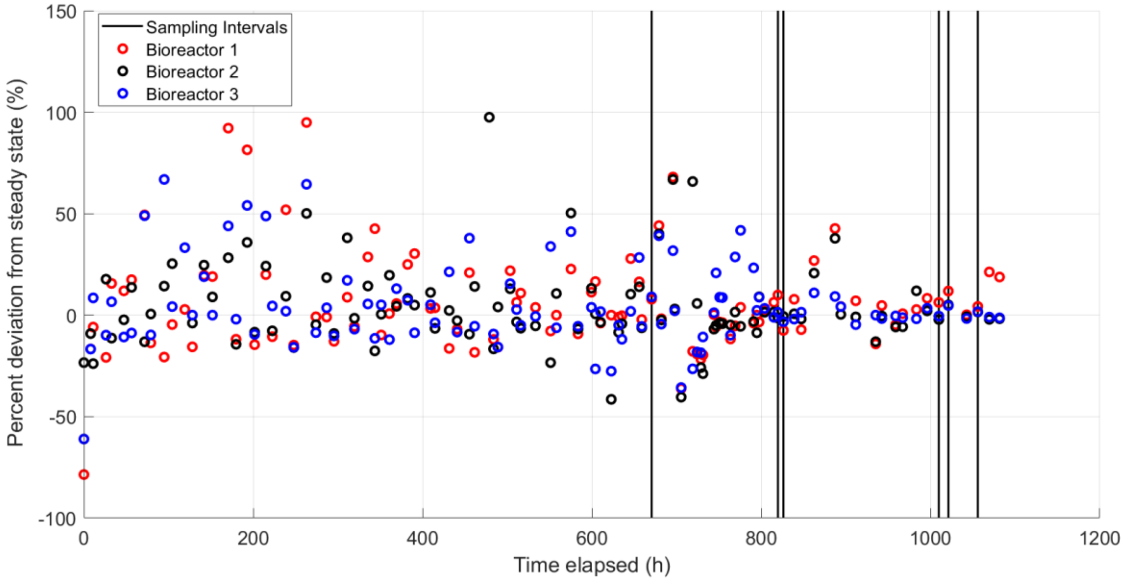


**Figure S7.** Time series of calculated deviations from theoretical steady-state for the 10h/18h/30h turnover time experiments. Samples were collected and analyzed on the condition that they represented a time interval of at least three complete turnovers at low deviation from steady state (< ±10%).

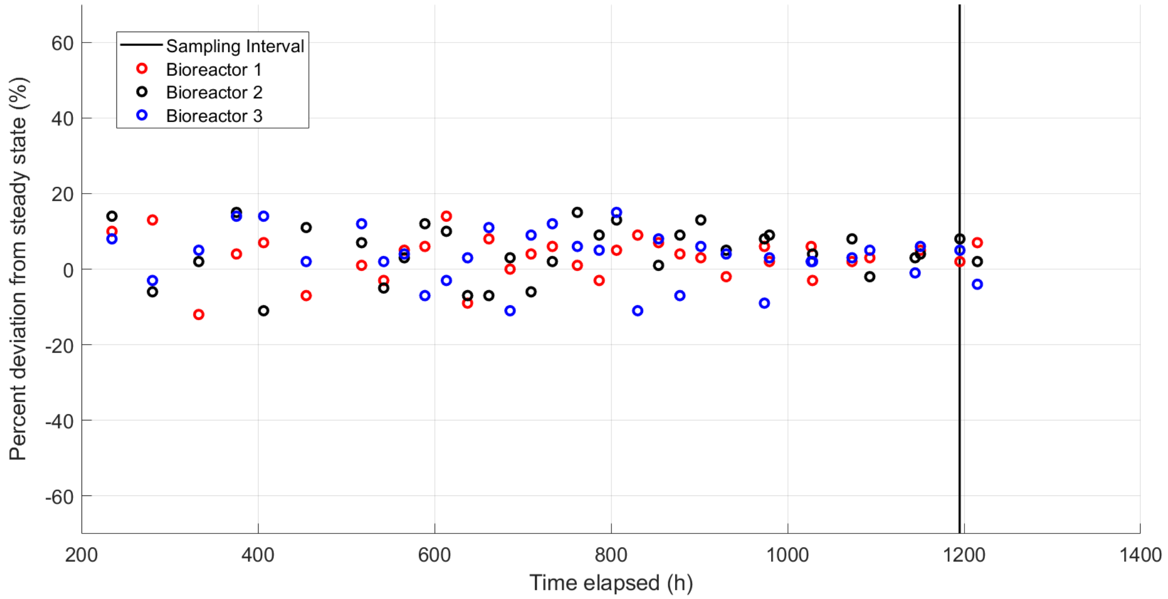


**Figure S8.** Time series of calculated deviations from theoretical steady-state for the 70h turnover time experiment. Samples were collected and analyzed on the condition that they represented a time interval of at least three complete turnovers at low deviation from steady state (< ±10%). Two types of samples were collected at the sampling interval denoted: One set of typical cold trap samples as well as one set of biomass pulled directly from bioreactors.

**Table S1**. Masses of freeze-dried cell pellets subjected to lipid extractions. Samples labeled 'Trap' correspond to a time-integrated volumes of cell culture collected in cold traps and spun down, with 50 mL of culture allocated to each pellet. Samples labeled 'Reactor' corresponds to pellets spun down from 10 mL volumes pulled directly from each reactor; these provide instantaneous "snapshots" of in situ lipid distributions. No significant difference in GDGT distributions was observed between these two sample types.

| **Sample ID** | **Freeze-dried pellet mass (g)** | **Sample Type** |
|---|---|---|
| S.aci_BR1S11_A_18h | 0.008 | Trap |
| S.aci_BR2S11_A_18h | 0.010 | Trap |
| S.aci_BR3S11_A_18h | 0.013 | Trap |
| S.aci_BR1S16_A_10h | 0.005 | Trap |
| S.aci_BR2S16_A_10h | 0.010 | Trap |
| S.aci_BR3S16_A_10h | 0.015 | Trap |
| S.aci_BR1S17_A_10h | 0.010 | Trap |
| S.aci_BR2S17_A_10h | 0.012 | Trap |
| S.aci_BR3S17_A_10h | 0.011 | Trap |
| S.aci_BR1S19_A_30h | 0.020 | Trap |
| S.aci_BR2S19_A_30h | 0.020 | Trap |
| S.aci_BR3S19_A_30h | 0.022 | Trap |
| S.aci_BR1S20_A_30h | 0.020 | Trap |
| S.aci_BR2S20_A_30h | 0.025 | Trap |
| S.aci_BR3S20_A_30h | 0.021 | Trap |
| S.aci_BR1S22_A_30h | 0.022 | Trap |
| S.aci_BR2S22_A_30h | 0.026 | Trap |
| S.aci_BR3S22_A_30h | 0.026 | Trap |
| Saci_BR2_S4_A_70h | 0.024 | Trap |
| Saci_BR3_S4_A_70h | 0.008 | Trap |
| Saci_BR1_S4_B_70h | 0.022 | Trap |
| Saci_BR3_S4_B_70h | 0.010 | Trap |
| Saci_BR2_S4_B_70h | 0.028 | Trap |
| Saci_BR1_S4_A_70h | 0.017 | Trap |
| Saci_BR1_S4_E_70h | 0.003 | Reactor |
| Saci_BR2_S4_F_70h | 0.003 | Reactor |
| Saci_BR1_S4_F_70h | 0.002 | Reactor |
| Saci_BR3_S4_E_70h | 0.003 | Reactor |
| Saci_BR3_S4_F_70h | 0.002 | Reactor |
| Saci_BR2_S4_E_70h | 0.003 | Reactor |

/end_file
